# The alignment of respiration to sensory-motor events is shaped by expected effort

**DOI:** 10.1101/2025.09.24.678231

**Authors:** Christoph Kayser, Lena Hehemann, Lisa Stetza

**Author notes:** Corresponding author: Christoph Kayser.

## Abstract

Humans often align their respiration with external events, a phenomenon thought to optimize neural resources for perception and action. Indeed, in sensory-cognitive experiments participants tend to align their respiration to the upcoming expected trials and their respiratory phase relates to neurophysiological processes reflecting changes in neural excitation, attention or arousal. However, it remains unclear whether this alignment is a passive entrainment to a task’s overall rhythm or an active process selectively aligning respiration based on the demands of individual events. We here tested this by recording respiration during three visual discrimination experiments that manipulated trial importance by either imposing different response deadlines or by manipulating trial value and difficulty. Our results show that participants align their respiration more consistently around stimulus onset for trials with short deadlines or trials presenting high-value and high-difficulty. These findings demonstrate that respiratory alignment is dynamically modulated on a trial-by-trial basis according to the anticipated required effort or task demands. Hence we conclude that respiration serves as an active tool to strategically allocate cognitive resources for sensory-motor challenges.

## Introduction

When preparing to perform specific actions or to act upon expected stimuli we often tend to temporally align our respiration with these events. In fact, the active regulation of respiration is fundamental in some sports or activities demanding precise manual coordination (McCarthy, 1995; Harbour et al., 2022). However, recent studies suggest that also during seemingly simple laboratory tasks probing sensation or cognition participants tend to structure their respiratory pattern around experimental events like stimulus presentation and response time (Huijbers et al., 2014; Nakamura et al., 2018; Grund et al., 2022; Johannknecht and Kayser, 2022; Goheen et al., 2024; Harting et al., 2025). This respiratory alignment explains some of the trial-to-trial variability in response accuracy and reaction times, suggesting that behavioral performance systematically comodulates with respiration (Flexman et al., 1974; Gallego et al., 1991; Vlemincx et al., 2011; Huijbers et al., 2014; Zelano et al., 2016; Arshamian et al., 2018; Melnychuk et al., 2018; Nakamura et al., 2018; Perl et al., 2019). This alignment of respiration may not only serve to control body movements (e.g. during shooting) but also to guide neural resources, such as excitability or arousal, to ensure the optimal processing of relevant sensory-motor contingencies, in line with theories of active sensing (Lakatos et al., 2019; Allen et al., 2023; Brændholt et al., 2023; Andrews et al., 2025).

Following this notion, recent studies have delineated the relation between respiration and brain activity in detail (Varga and Heck, 2017; Brændholt et al., 2023; Engelen et al., 2023; Kluger et al., 2024; Madsen and Parra, 2024). The brain structures controlling respiration and those sensing the resulting changes in airflow or chest pressure are intricately connected with the limbic system (Del Negro et al., 2018; Ashhad et al., 2022; Krohn et al., 2023). As a result, direct neural feedback about the respiratory state is widely available in the brain (Heck et al., 2016; Tort et al., 2018; Kluger and Gross, 2021). Critically, both aperiodic signatures of neural excitability as well as rhythmic signatures of arousal seem to comodulate with the respiratory phase (Kluger et al., 2021; Kluger et al., 2023) and respiration directly modulates the encoding of task-specific sensory signals in relevant brain regions (Park and Tallon-Baudry, 2014; Zelano et al., 2016; Perl et al., 2019; Stetza et al., 2025). This suggests that by actively aligning respiration with expected stimuli or actions our brain (consciously or unconsciously) ensures an optimal neural substrate to process or perform these.

Building on these findings, we previously investigated the relation between respiration and task performance across multiple datasets comprising perceptual and cognitive tasks (Harting et al., 2025). We demonstrated that both the trial-wise respiratory phase preceding a trial and the trial-averaged respiratory phase of individual participants are predictive for their behavioral outcomes: participants who aligned their respiration ‘optimally’ to the paradigm tended to respond faster than those how aligned ‘non-optimally’ (Harting et al., 2025). If respiration indeed functions as a mechanism to align neural resources to important events, one could assume that this alignment of respiration is particularly strong when events are expected to place higher demands or to require high effort to be completed successfully. Under the assumption that the nervous system attributes it limited resources optimally (Barlow, 1961; Lennie, 2003), one would hence expect stronger alignment to more important or effort-demanding events and a reduced alignment to less important or less effort-demanding events.

Yet, most previous studies on respiration-task alignment only focused on the overall alignment of respiration regardless of task demands and did not explicitly manipulate the incentive to optimize the alignment of respiration to individual trials by manipulating trial importance (Huijbers et al., 2014; Nakamura et al., 2018; Grund et al., 2022; Johannknecht and Kayser, 2022; Goheen et al., 2024; Harting et al., 2025). Hence it remains unclear whether the hypothesis of a differential alignment to more and less important events indeed holds true. We here test this hypothesis based on three perceptual tasks in which we manipulated trial-wise importance, operationalized either by the requirements for response speed (by imposing loose or tight response deadlines) or by the explicit manipulation of trial value and difficulty. Overall, our data support that participants align their respiration more reliably to events with higher demands or importance.

## Methods

### Participants and general procedures

The study was approved by the ethics committee of Bielefeld University. Adult volunteers with self-reported normal vision and hearing participated after providing informed consent. All participants were compensated for their time. The data were collected anonymously and it is possible that some individuals participated in more than one of the experiments described below. Demographical data were not collected and the participant pool consisted of typical young university students. Prior to the study, our interest to investigate the relation between respiration and task performance was not mentioned explicitly and participants were instructed to “breathe through their nose as usual”, as if performing the experiments without wearing a mask. During the experiment we could not continuously monitor whether participants adhered to this instruction, leaving the possibility that during parts of the experiment participants were breathing orally.

The experiments were performed in a darkened and sound-proof booth (E: Box; Desone, Germany). Visual stimuli were presented on a computer monitor (27” monitor; ASUS PG279Q, about 85 cm from participant’s head) with stimulus presentation controlled using the Psychophysics Toolbox (Version 3.0.14) in MATLAB (Version R2017a; The MathWorks, Inc., Natick, MA). Participants responded using a computer keyboard.

As in our previous studies, respiration data were recorded using a temperature-sensitive resistor (Littelfuse Thermistor No. GT102B1K, Mouser electronics) that was inserted into disposable clinical masks, capturing the temperature changes resulting from the respiration-related airflow (Johannknecht and Kayser, 2022; Harting et al., 2025). The voltage drop across the thermistor was recorded via an ActiveTwo EEG system (BioSemi BV) at a sampling rate of or 1000 Hz.

### Behavioral paradigms

We collected data in three experiments probing visual tasks that were also used in previous studies on respiration (Johannknecht and Kayser, 2022; Harting et al., 2025). Experiment 1, which manipulated response deadlines, relied on an emotion discrimination paradigm; experiments 2 and 3, which manipulated task difficulty and trial-value, relied on a random dot motion discrimination paradigm (Fig. 1A). All experiments involved a practice block, for experiments 2 and 3 a block to determine each participant’s perceptual threshold for the respective dot motion task and one or more blocks of experimental trials. For each task, participants were instructed to respond as fast and accurately as possible after stimulus onset.

**Figure 1.**
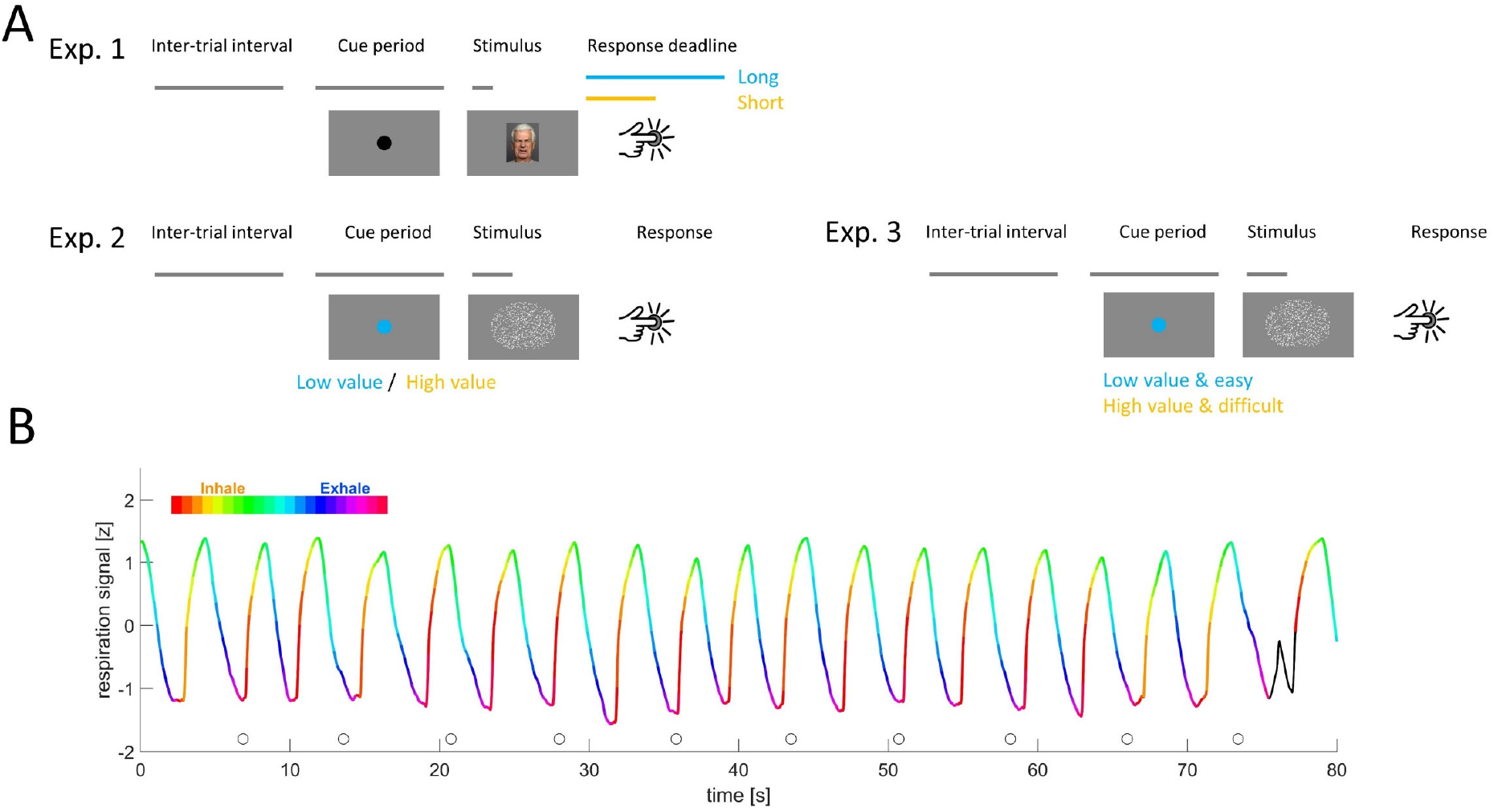
Paradigms and example data. **A)** Schematic of the three experiments. Each experimented featured pseudo-random inter-trial intervals, a fixed cueing period indicating the upcoming stimulus, a stimulus and a response period. Experiment 1 featured an emotion discrimination task and contrasted conditions with long and short response deadlines presented in different blocks. Experiment 2 and 3 featured a random dot motion task and manipulated trial value and difficulty. Experiment 2 manipulated both independently, while experimented 3 yoked them. Trial value was indicated by the cue. B) Example respiratory trace from one participant. Color indicates the respiratory phase, inhalation is shown upwards. Circles indicate stimulus onset times. The black part of the curve is an example of an atypical cycle.

In contrast to our previous studies using these paradigms, we here presented trials with longer inter-trial intervals and with temporally reliable cues that indicated the expected stimulus onset. In previous studies, the pre-stimulus (fixation) periods were pseudorandom and on the order of 400-1000ms. Hence they provided only somewhat reliable temporal cues about stimulus onset. We here relied on longer but fixed fixation periods (see below) to allow the precise expectation of when a stimulus would appear. We also included longer inter-trial intervals compared to previous work. In our previous studies, the inter-trial intervals were on the order of 1.3 seconds, resulting in an overall pacing of subsequent trials on the order of around 3 seconds, as typical for behavioral and neuroimaging paradigms probing sensory or cognitive processes (see Fig. 1B. in(Harting et al., 2025)). The design implemented here resulted in time intervals between subsequent stimulus onsets of 5.75±0.10 s (mean±SD) for experiment 1 and 7.58±0.36 s and 7.42±0.16 for experiments 2 and 3. This time scale is longer than the typical respiratory cycle duration for most participants. These changes were introduced to allow a better alignment of respiration to the experimental trials based on temporal expectation by allowing for longer inter-trial intervals commensurate with respiration and by providing clear predictive cues.

*Experiment 1:* Participants discriminated the emotional expression (sad or happy) of briefly presented faces (subtending 15 x 12 degrees, presented for ∼17 ms). Stimuli were obtained from the Dynamic FACES database (Holland et al., 2019) and each trial presented a different image. Trials started with a fixation period indicated by the appearance of a central fixation dot that remained on the screen until stimulus onset. This pre-stimulus period lasted 1000ms and was fixed. Inter-trial intervals were drawn from a uniform distribution between 3500-4000ms. Importantly, this task was performed either in the context of a LONG response deadline, or a SHORT response deadline. In the LONG condition participants had 1.4 s time to submit their responses, in the SHORT condition they were given a reaction time (RT) deadline that was titrated for each participant based on their actual RTs in the LONG condition. The threshold was defined as the 30% percentile of the RT distribution in the LONG blocks. Participants performed four blocks of 100 trials for each condition, with the first half of the experiment presenting the LONG condition and the second half the short condition (hence 800 trials in total). If participants did not submit a response within a deadline period they received visual feedback and those trials were excluded for analysis. Overall we obtained data from n=27 participants, for which we retained 643 ± 19 trials on average (mean ± s.e.m.) for the analysis.

*Experiment 2:* Participants judged the direction of motion (left- or right-wards) of visual random dot displays. Random dot displays lasted 660 ms, subtended 8 degrees of visual angle and contained 800 limited-lifetime dots (0.2° diameter, 8 frames life-time) moving at 3.5 degrees per second. The coherence of dots (fraction of dots moving in the same direction) manipulated task difficulty. Trials could either be easy or hard, with hard trials presenting motion coherence around participants perceptual threshold and the easy condition featuring a coherence that was 3.5% higher than threshold. Prior to the actual task we determined participants perceptual thresholds (around 72% correct responses) using three interleaved one-up two-down staircases with multiplicative step-sizes. Importantly, within the experimental blocks each trial was assigned either a high or a low value. Value was manipulated independently of task difficulty, and both value and difficulty were pseudo-randomized across trials. Participants were told that they could earn points during experiment for submitting correct responses and that there were trials with higher value (yielding 3 points) and with lower value (1 point). High value trials were less frequent (relative ratios of high and low value 1:2.33). At the end of the experiment participants were compensated for their time, but also received additional compensation based on their score (computed as their total score / 100). Participants performed 5 blocks of 132 trials (660 trials in total). For participants to be able to exploit the value of each trial they were presented with cue prior to each stimulus presentation: this was a 3-sec counter, counting down the seconds until stimulus onset. The color of this indicated the value of the upcoming trial. Inter-trial intervals were 1700-2800ms (uniform). We obtained data from n=24 participants for which we retained 585 ± 14 trials on average

*Experiment 3:* This experiment featured the same overall design as experiment 2. But rather than value and difficulty being independent variables, these were linked. That is, the high value trials also had higher difficulty, and the low value trials had lower difficulty. In addition, we increased the coherence difference between easy and difficult trials, presenting the high value trials around threshold and the low value (and easier) trials at 8 % above perceptual threshold.). Participants performed 5 blocks of 126 trials (630 trials in total). We obtained data from n=23 participants for which we retained 601 ± 12 trials on average.

### Data preprocessing

The respiratory data were separated into individual cycles as described previously, based on the Hilbert transform and the detection of local peaks and throughs (Harting et al., 2025). We characterized atypical respiratory cycles by comparing the time course of individual cycles using their mean-squared distances. For each participant we excluded cycles with a distance larger than 3 standard deviations from the centroid distribution as atypical, as these may reflect sights of breath holds. Within each cycle we defined the respiratory phase as a linearly increasing variable from the beginning to the end of inspiration (defined as angle from 0 to pi) and subsequently as linearly increasing from the beginning to the end of expiration (defined as pi to 2*pi). This phase variable was resampled to 20Hz for subsequent analysis. From the overall data we removed trials for which the respiratory cycle at the time of stimulus onset was atypical and trials with reaction times shorter than 200 ms or longer than 3 s. For experiment 1 we also discarded trials without responses within the required deadline. Respiratory example data from one participant are shown in Figure 1B.

### Data analysis - alignment of respiration

As in previous work we quantified the alignment of respiration to the experimental paradigm using a measure of phase-locking vector strength (plvs). This was obtained by first defining the respiratory phase as a complex-valued number. Then we averaged these complex numbers across trials and used the vector length as plvs (Kluger et al., 2021; Johannknecht and Kayser, 2022). To contrast the actual plvs against the null hypothesis of no alignment, we derived a surrogate distribution of plvs for each participant by randomly time-shifting the respiratory trace and recalculating the phase consistency 4000 times (Harting et al., 2025). We then compared the actual group-mean to the distribution of group-means in the surrogate data, taking the maximal value along the time axis to correct for multiple comparisons.

Our main hypothesis concerned whether this alignment differs between conditions placing different demands or importance. For this we compared the group-level plvs between conditions using paired t-tests and relied on a cluster-based permutation procedure correcting for multiple comparisons over time (Maris and Oostenveld, 2007). For this we shuffled the condition-difference independently for each participant and obtained the distribution of differences under the null hypothesis of no difference between conditions across 4000 shuffles. We then probed for significant clusters in the actual data based on a minimal cluster size of 250 ms and using the max-sum for cluster-forming. For these tests we report p-values and the Cohen’s D effect size at the time point of largest difference within the cluster.

As a second measure of respiratory dynamics we quantified the slope by which the plvs changes around the trial onset. For this we computed the difference between the minimal value in the time period of -2 to -1 s prior to stimulus onset to the maximal value in the window of +1 to +2 s. This slope reflects how quickly any change in respiratory alignment unfolds around stimulus onset. We contrasted these slopes between conditions using paired t-tests. For these we report p- and t-values and Cohen’s D as effect size.

### Data analysis - respiratory cycle durations

For each trial we determined the respiratory cycle including stimulus onset time and calculated the duration of this cycle. We then contrasted the trial-averaged durations between conditions at the group-level using paired t-tests. In addition, we correlated the participant-wise condition-difference in cycle duration with the condition-difference in response accuracy and reaction times using Spearman’s rank correlation. The respective r- and p-values are indicated in the respective scatter plots.

## Results

We tested the active alignment of respiration to the sequence of trials in three experiments manipulating either the requirement on response speed (experiment 1) or the importance of submitting a correct response by manipulating trial value (experiment 2 and 3). The experimental tasks were taken from previous studies demonstrating that participants tend to align their respiration to experimental trials also in the presence of temporally imprecise cues for stimulus onsets. We here adjusted the experimental design to feature longer inter-trial intervals and fixed pre-stimulus periods to allow the precise expectation of when a stimulus would appear. In the present data the actual time intervals between subsequent stimulus onsets were on the order of 5.7 to 7.5 seconds (see Methods) This time scale is longer than the typical respiratory cycle duration for most participants and hence should be compatible with an active alignment of respiration to the experimental sequence. For the present data the average duration of respiratory cycles across all three experiments was 3.79 ± 0.93 s (mean ± SD).

### Influence of respond deadline

We first asked how changes in the demands on response speed would affect the respiratory alignment to the experimental trials. In experiment 1 we contrasted trials performed with a LONG reaction time deadline (1.4 s) to trials performed on a SHORT deadline (tailored to the 30^th^ percentile of RTs observed during the LONG condition). As expected, reaction times were significantly shorter in the SHORT compared to the LONG condition (Fig. 2A; 0.461 ± 0.013 s vs. 0.618 ± 0.021 s, paired t-test p<10^-5^, t=16.17, Cohen’s D = 3.17) and response accuracy was significantly lower in the SHORT condition (90.7 ± 1 % vs 96.8 ± 0.3 % correct responses, p<10^−5^, t=7.77, Cohen’s D = 1.52).

**Figure 2.**
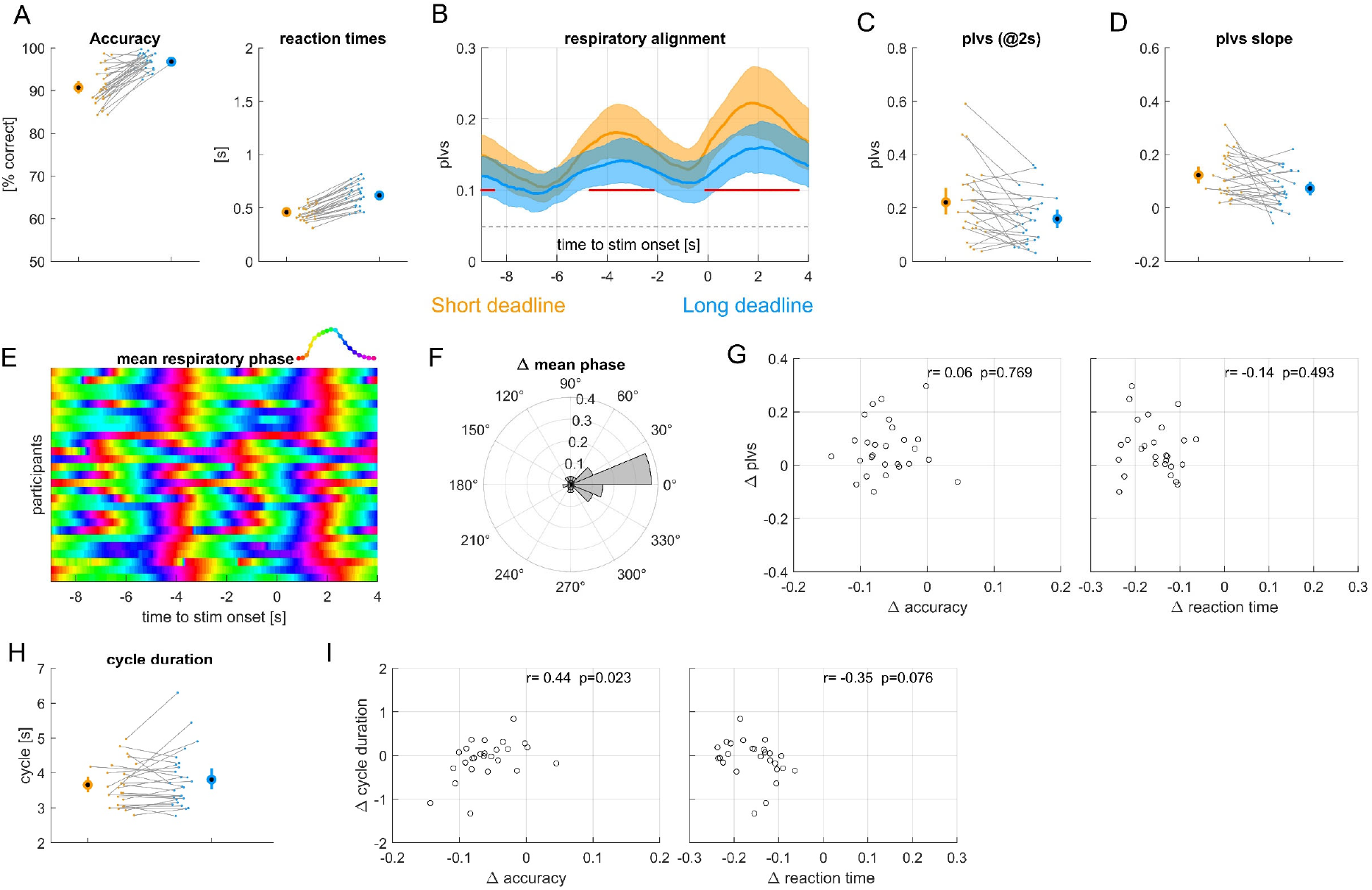
Results from experiment 1 manipulating response deadlines. **A)** Accuracy and reaction times for short and long response deadlines (see color code). **B)** Alignment of respiration to stimulus onset, measured using the phase-locking vector strength (plvs). Red lines indicate significant differences (cluster-based permutation test, at p<0.01). **C)** Participant- and condition-wise plvs values at +2s. **D)** Slope of plvs change around stimulus onset. **E)** Trial-averaged respiratory phase for each participant, greenish colors indicate peak inhalation. **F)** Condition difference in trial-averaged respiratory phase shown as angular histogram. **G)** Participant-wise condition-differences in plvs vs. the differences in response accuracy and reaction times. r- and p-values are based on a Spearman rank correlation. **H)** Average duration of those respiratory cycles including stimulus onset. **I)** Participant-wise differences in respiratory cycle duration vs. differences in response accuracy and reaction times. Group-level data are shown as mean and 95% percentile bootstrap confidence intervals.

The within-participant alignment of the respiratory phase was higher around the stimulus-response times compared to the period a few seconds earlier, as expected based on previous work. This phase locking was significantly stronger than expected by chance (permutation test against surrogate data; p<0.01 corrected for multiple tests along time). More importantly, this phase locking was significantly stronger in the SHORT compared to the LONG condition (Fig. 2B). A cluster-based permutation test correcting for multiple comparisons over time revealed two significant clusters (cluster 1: -4.65 to - 2.25 s, p=0.004, Cohen’s D = 0.58; cluster 2: -0.10 to 3.55 s, p=0.004, Cohen’s D = 0.74). Figure 2C illustrates the participant wise plvs values for both conditions at t=2s. To further characterize this dynamic alignment of respiration we computed the slope of the plvs around each stimulus onset (Fig. 2D). This slope reflects how quickly respiratory alignment synchronizes to the expected stimulus onset and was significantly higher in the SHORT compared to the LONG condition (p=0.0087, t=2.84, Cohen’s D = 0.55). Hence, our data show that participants tended to align their respiratory phase to the expected upcoming trials significantly tighter in the SHORT condition.

To probe how similar or different individual participants tended to breath around trials we compared the trial-averaged respiratory phases between participants (Fig. 2E). Visibly, this average phase was comparable across participants and most participants were at peak inhalation around stimulus onsets. Importantly, across participants the trial-average respiratory phase was also similar between SHORT and LONG conditions (Fig. 2F; randomization test comparing the mean angles, mean difference 0.101 radians, 95% percentile CI: [-0.378, 0.353], p=0.563).

We next asked whether the condition difference in the behavioral data relates to the condition difference in plvs. For this we computed the correlation of the condition difference in plvs to the respective differences in reaction times and accuracy across participants. These correlations were not significant (Fig. 2G; see Spearman rank correlations indicated there).

Finally, we probed whether the durations of participants respiratory cycles differed between the two conditions. For this we extracted for each trial the duration of that respiratory cycle including stimulus onset. These durations did not differ significantly between conditions (Fig. 2H; paired t-test, p=0.184 t=-1.37, Cohen’s D = 0.26). However, the within participant condition differences did correlate significantly with the condition effects on response accuracy (Fig. 2I; r=0.44, p=0.023) and were close to significance for reaction times (r=-0.35, p=0.076). The data suggest that those participants whose behavior was least affected by the SHORT reaction time deadline were those who exhibited comparable respiratory cycle durations in both conditions or who even tended to show longer respiratory cycles during the SHORT condition.

### Influence of trial value independent of task difficulty

In a second experiment we manipulated task difficulty and the trial-wise value of correct responses as independent factors. We expected that participants would structure their respiratory phase more systematically around high compared to low value trials, and that they would also perform better in these trials. Yet, the behavioral data revealed only very small effects of stimulus level and value. For response accuracy neither effect was significant (Fig. 3A; value: F=0.62, p=0.433; level: F=0.24, p=0.628) and the numerical difference between high hand low value trials was minimal (high value: 80.5 ± 1.5%, low value: 80.0 ± 1.4%). For reaction times the effect of value but not of level was significant (value: F=11.4, p=0.001; level: F=0.07, p=0.794), and reaction times were longer for high value trials (high value: 1.25 ±0.05 s, low value: 1.233 ± 0.04 s), although numerically the effect was small. Hence, the manipulations of value and difficulty in this experiment had only a minimal influence on behavior, suggesting that any effect of value on respiration may also be small.

**Figure 3.**
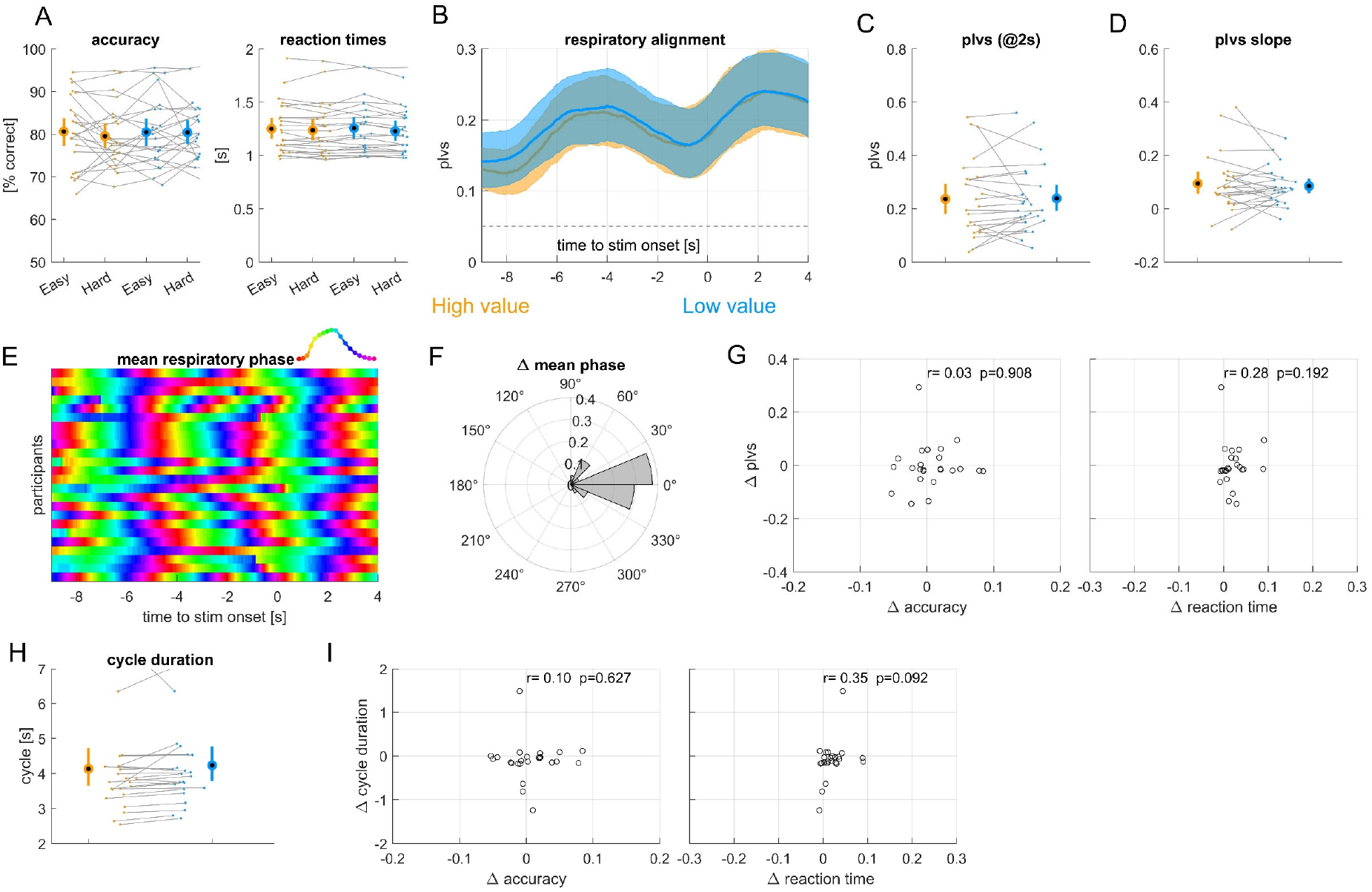
Results from experiment 2 manipulating trial value and difficulty independently. **A)** Accuracy and reaction times for each condition. High value trials are orange, low value trials blue. Behavioral data are additionally split by difficulty (easy, hard). **B)** Alignment of respiration to stimulus onset, measured using the phase-locking vector strength (plvs) for each value, averaged over difficulty levels. **C)** Participant- and condition-wise plvs values at +2s. **D)** Slope of plvs change around stimulus onset. **E)** Trial-averaged respiratory phase for each participant, greenish colors indicate peak inhalation. **F)** Condition difference in trial-averaged respiratory phase shown as angular histogram. **G)** Participant-wise condition-differences in plvs vs. the differences in response accuracy and reaction times. r- and p-values are based on a Spearman rank correlation. **H)** Average duration of those respiratory cycles including stimulus onset. **I)** Participant-wise differences in respiratory cycle duration vs. differences in response accuracy and reaction times. Group-level data are shown as mean and 95% percentile bootstrap confidence intervals.

Indeed, while the overall alignment of respiration the experimental paradigm was significant, the plvs did not differ between high and low value conditions (Fig. 3B; a cluster-based permutation test did not reveal any significant clusters at p<0.05). The slope of the phase locking values also did not differ significantly (Fig. 3D, paired t-tests, p=0.522, t=0.65, Cohen’s D =0.13). As for experiment 1 participants tended to reach peak inhalation at the time of stimulus onset (Fig. 3E) and the trial-averaged respiratory phase did not differ between value conditions (angular difference: mean = 0.10 radians, 95% CI: [-0.211, 0.195], p=0.369, Fig. 3F). The correlations between condition differences in behavior and respiration were not significant (Fig. 3G,I; see correlation values there) and the duration of respiratory cycles was also comparable between conditions (Fig. 3H; p=0.285, t=-1.09, Cohen’s D =-0.223).

### Influence of lined trial value independent and task difficulty

The lack of effects in experiment 2 may have resulted from an overall low effectiveness of the manipulation of value, and or the fact that high and low value tasks had on average comparable difficulty, and hence possibly required similar efforts. We hence repeated this experiment but this time linking task difficulty and value, rending the high value trials more difficult compared to the low value trials. In addition, we increased the difference in task difficulty between conditions. This combined manipulation of value and difficulty resulted in a significant difference in reaction times between conditions, with reaction times in high value trials being longer compared to low value trials (Fig. 4A; high value: 1.158 ± 0.033 s; low value: 1.140 ± 0.030; p=0.040, t=2.18, Cohen’s D = 0.46). Response accuracy did not differ significantly between conditions (high value: 74.3 ± 1.4 %, low value: 75.7 ± 1.7 %; p=0.110, t=-1.66, Cohen’s D = 0.35).

**Figure 4.**
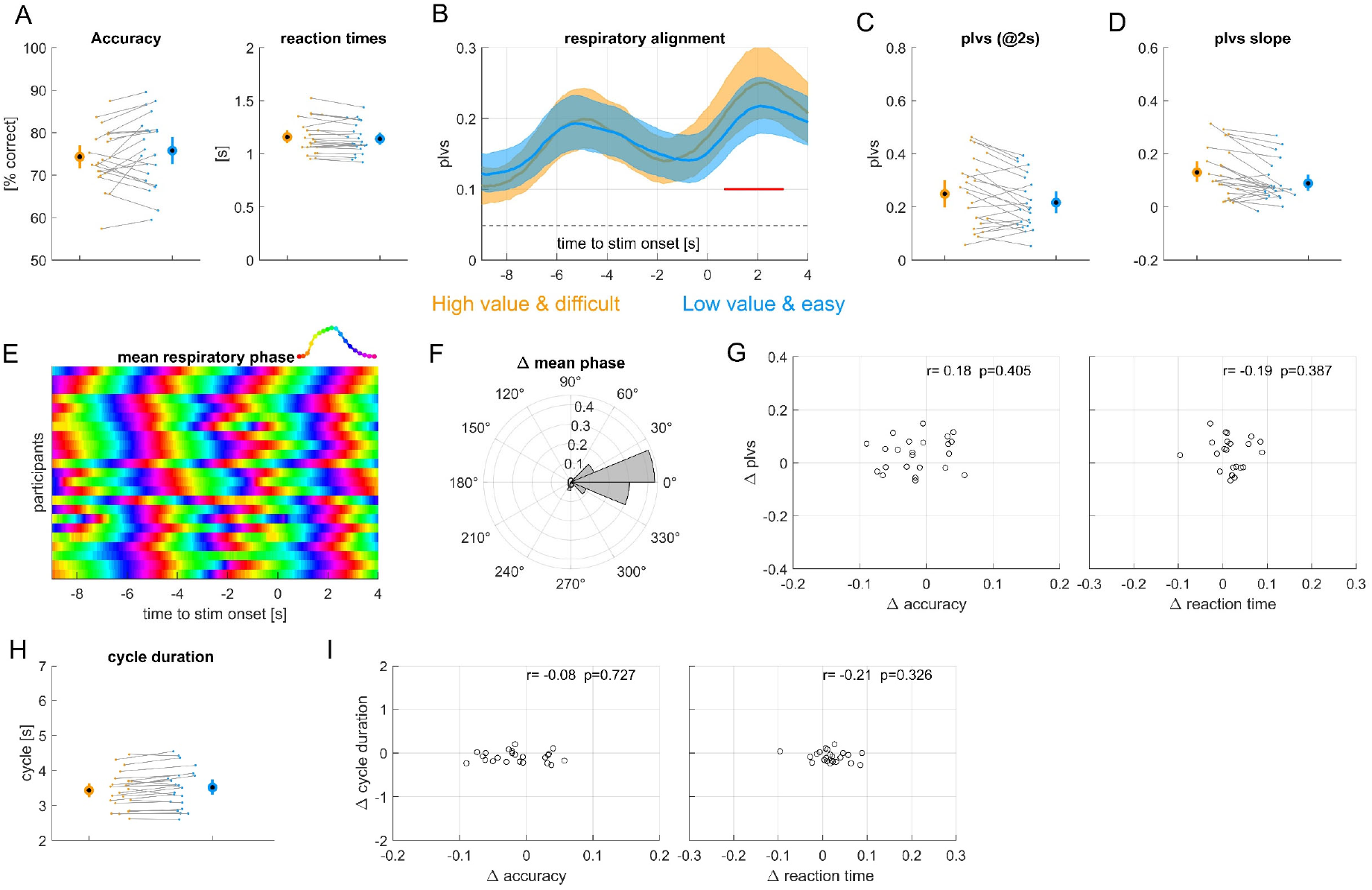
Results from experiment 3 linking trial value and difficulty. **A)** Accuracy and reaction times for high value & difficult trials (orange) and low value & easy trials (blue). **B)** Alignment of respiration to stimulus onset, measured using the phase-locking vector strength (plvs). Red lines indicate significant differences (cluster-based permutation test, at p<0.01). **C)** Participant- and condition-wise plvs values at +2s. **D)** Slope of plvs change around stimulus onset. **E)** Trial-averaged respiratory phase for each participant, greenish colors indicate peak inhalation. **F)** Condition difference in trial-averaged respiratory phase shown as angular histogram. **G)** Participant-wise condition-differences in plvs vs. the differences in response accuracy and reaction times. r- and p-values are based on a Spearman rank correlation. **H)** Average duration of those respiratory cycles including stimulus onset. **I)** Participant-wise differences in respiratory cycle duration vs. differences in response accuracy and reaction times. Group-level data are shown as mean and 95% percentile bootstrap confidence intervals.

The within-participant phase locking of respiration was significant, and importantly, was significantly stronger in the high-compared to the low value condition (cluster-based permutation test revealed a significant cluster from 0.7 to 3.0 s, p=0.004, Cohen’s D = 0.55; Fig. 4B and the individual data in Fig. 4C). Furthermore, this difference in plvs was accompanied by a difference on phase locking dynamics, as the slope of plvs was significantly higher in the high value condition (Fig. 4D; p=0.001, t=3.64, Cohen’ D = 0.76). Similar to the other experiments, the trial-averaged phase angles did not differ systematically across participants (Fig. 4E,F; angular difference: mean = -0.05 radians, 95% CI: [-0.180, 0.184], p=0.685) and the condition-differences in plvs did not correlate significantly with the respective differences in accuracy or reaction times (Fig. 4G).

The durations of respiratory cycles differed significantly between conditions (p=0.004, t=-3.22, Cohen’s D =0.67; Fig. 4H), with shorter cycles in the high value condition. However, these condition differences in respiratory cycle duration did not correlate with differences in accuracy or reaction times (Fig. 4I). Overall, participants aligned their respiration more strongly to the high-value (and difficult) trials and did so with shorter breathing cycles. Taken together, across experiments 2 and 3, respiration showed robust alignment to task events, but this alignment was only modulated when task value was linked with difficulty.

## Discussion

Previous studies have shown that participants tend to align their respiratory cycles to expected (albeit sometimes pseudo-randomly timed) experimental trials (Huijbers et al., 2014; Nakamura et al., 2018; Grund et al., 2022; Johannknecht and Kayser, 2022; Goheen et al., 2024; Harting et al., 2025). The general assumption is that this adaptation leads to an advantage in behavior and perception, as the brain is in an optimal state to process a stimulus. However, such alignment could also reflect synchronization to the overall experimental pace rather than to trial-specific demands, such as effort or difficulty. The hypothesis that respiration is actively structured to support skilled actions or optimize sensory perception implies that, during moments of high behavioral significance, aligning respiration more closely with task events may confer an advantage. We here probed this hypotheses by manipulating the required effort by either imposing short response deadlines or manipulating trial difficulty. Overall our data support that participants tend to breathe more consistently on trials requiring more effort to respond accurately and support respiration as an active tool to guide neural resources to challenging sensory-motor events (Allen et al., 2023; Brændholt et al., 2023; Engelen et al., 2023; Kluger et al., 2024; Andrews et al., 2025).

### The alignment of respiration to critical moments

It is known that humans tend to breath in a structured manner around expected events, such as during sports, during conversation and also in a laboratory task (Rochet-Capellan and Fuchs, 2014; Palumbo et al., 2017; Harbour et al., 2022; Johannknecht and Kayser, 2022; Goheen et al., 2024). This alignment of respiration to such events manifests in a consistency of the respiratory phase across trials. In previous experiments, the timing of experiment trials was often pseudo-random, as is common for neuro-cognitive laboratory tasks. This temporal uncertainty is often on the scale of about 200-1000ms, and introduced to avoid effects of precise temporal expectation, which can engage mechanisms of temporal prediction and can act as a confounder depending on the experimental question (Nobre et al., 2007; Schroeder and Lakatos, 2009; Nobre and van Ede, 2023). Despite this temporal uncertainty participants tend to structure their respiration around the experimental events, which typically unfold on a time scale of several seconds, a scale similar to that of individual respiratory cycles. Hence, it could be that in these previous studies participants aligned their respiration to the overall time scale of the experiment, rather than to the individual stimuli or motor actions.

Yet, the notion that the brain optimally allocates neural resources in general predicts that these resources should be particularly deployed for important events, i.e. those that require greater effort or which otherwise hold particular value. Recent studies suggest respiration serves to structure neurophysiological processes related to attention, arousal and neural excitability (Del Negro et al., 2018; Tort et al., 2018; Kluger et al., 2021; Kluger et al., 2023; Kluger et al., 2024; Stetza et al., 2025). Hence, specifically structuring respiration around critical events would set the brain into a state to optimally react to these. Our data support this hypothesis based on two experimental manipulations: one based on the combined difficulty and value of individual trials and one based on a manipulation of response deadlines. The latter required participants to exploit the sensory information more swiftly to respond, and punished late responses by not letting participants respond to this trial anymore. In the joint manipulation of value and difficulty, participants could earn more points on difficult compared to easy trials. In both they aligned their respiratory phase more reliably to trials requiring more effort to succeed.

In experiment 2, which manipulated trial value and task difficulty independently, we did not observe a significant effect of value on either behavior or respiration. One possible explanation could be the following: in this design only value was known in advance, whereas the difficulty could only be judged immediately during the stimulus presentation. Consequently, an optimal strategy for this task design may involve deploying only some effort towards adjusting neural resources according to the expected value and some effort towards adaptively deploying resources once difficulty is known (i.e. during stimulus). The division of resources could lead to weaker respiratory alignment compared to the alignment in a task including a joint manipulation of difficulty and value, which is consistent with our observations. Together our data suggest that the alignment of respiration is driven by the anticipated effort required to successfully complete the trial.

### Is respiratory alignment driven by sensory expectations or motor actions?

One yet unresolved question concerns whether the alignment of respiration to the experiment trials is driven by the temporal expectation of a specific stimulus, by the expectation to perform a specific action or the action itself (Harting et al., 2025). In principle, respiratory alignment could simply be driven by the expectation of something to happen, i.e. a stimulus to appear. However, in line with the notion of active sampling, we propose that this alignment would be stronger in the context of specific task requirements, hence the need for a motor response. On the time scales of the experiments performed here or in previous work these three factors are all highly correlated. The stimulus-response delays are short compared to the duration of respiratory cycles, and all trials required an active response. Hence, the current data cannot dissociate these factors. To better dissociate the potential drivers of the alignment, we need experiments with clearly expected and unexpected stimuli, as well as manipulations of the stimulus-response contingencies. One possibility would be to use a go/no-go paradigm, where a stimulus is reliably expected on every trial but a motor response is required on only a subset of them.

### Conclusion

The outcomes of perception and action are intricately linked to the physiological state of our body. Recent studies have shown that under laboratory conditions participants tend to structure their respiration systematically around expected events such as individual experimental trials. We here show that the strength of this alignment is specific to the value attributed to individual trials or the expected effort required. Hence, we tend to actively structure our respiration in a very selective manner for individual upcoming required actions.

## Acknowledgements

We would like to thank Sepideh Mirzaei for help with data collection.

